# eRNAkit: Expanding the Functional Atlas of human Enhancer RNAs Beyond the Nucleus

**DOI:** 10.1101/2025.04.25.650683

**Authors:** Natalia Benova, Rene Kuklinkova, Mahmoud K. Eldahshoury, Chinedu A. Anene

## Abstract

Enhancer RNAs (eRNAs) are a class of non-coding RNAs transcribed from active enhancers that regulate various aspects of transcription. Although traditionally viewed as nuclear-localised and unstable transcripts, large transcriptomic studies reveal they vary widely in localisation and biochemical properties, including detectable accumulation in cytoplasmic compartments. These observations suggest potential non-nuclear functions for eRNAs, yet existing databases remain focused on nuclear, cis-regulatory roles, limiting systematic exploration of their broader regulatory repertoire. Here we present **eRNAkit**, a comprehensive and accessible resource designed to address this gap by integrating subcellular localisation, RNA-RNA interaction and expression data for annotated eRNAs. Leveraging fractionation-based RNA-Seq datasets, eRNAkit profiles eRNA distribution across nuclear and cytoplasmic compartments. It incorporates gene expression data spanning major human organs and primary cell types, enabling tissue-specific analysis of eRNA function. Crucially, eRNAkit includes experimentally derived RNA-RNA interaction data from RIC-Seq, PARIS, and KARR-Seq, supporting exploration of trans-acting and cytoplasmic roles for eRNAs grounded in physical interaction evidence. eRNAkit expands current eRNA resources beyond the enhancer-promoter paradigm, offering a robust platform for dissecting non-canonical functions of eRNAs and advancing our understanding of their full regulatory potential in human biology. The eRNAkit resource is available to download at https://github.com/AneneLab/eRNAkit.

## Introduction

eRNAs are a class of non-coding RNAs transcribed from active enhancer regions ^1^. They regulate gene expression by activating transcription, bridging enhancer-promoter interactions, and remodelling chromatin ^2^. Although initially considered nuclear, unstable and non-polyadenylated, transcriptomic data — including those from the ENCODE consortium, reveal that eRNAs vary widely in both biochemical properties and cellular localisation ^3,4^. While most studies report nuclear enrichment based on relative abundance, they also detect eRNAs in cytoplasmic fractions, raising the question of what functions these cytoplasmic eRNAs might perform — and which molecular targets they may regulate outside the nucleus.

Existing eRNA databases predominantly emphasise *cis-*regulatory mechanisms, reflecting the prevailing model in which enhancers influence the expression of nearby genes through spatial proximity. Resources such as eRNAbase, HeRA, Anima-eRNAdb, eRic and GPIeR ^5–9^ systematically catalogue eRNA-target relationships based on genomic co-localisation, chromatin conformation, or correlative expression patterns. While these approaches have illuminated many aspects of enhancer biology, their design inherently favours local interactions and overlook more distal, cytoplasmic effects. Furthermore, these databases largely ignore the localisation of eRNAs, an essential factor in understanding their functional potential. Nevertheless, mounting evidence suggests that certain eRNAs can function beyond their sites of origin, modulating the expression of distant genes or engaging in post-transcriptional regulation. Some studies have uncovered eRNAs that act in *trans*, regulating gene expression far from their sites of origin. For example, a distal regulatory region eRNA transcribed from the MYOD1 locus mediates cohesion recruitment and Myogenin gene expression in trans to control myogenic differentiation ^10^. Similarly, an eRNA transcribed from the KLK3 locus enhance androgen receptor-dependent gene expression in human prostate cancer ^11^. In both examples, the eRNAs were found to be polyadenylated, which is contrary to the standard model of eRNA processing. In a different study, knockdown of two p53-induced enhancer RNAs, p53BER2 and p53BER4, impaired the induction of distant target genes in response to p53 activation ^12^. This regulation occurred despite the absence of detectable p53 binding at the promoters of these target genes, suggesting that the eRNAs act independently of local enhancer–promoter contact. Moreover, given that siRNA knockdowns used to validate most of these observations are more effective for cytoplasm localised RNAs ^13^, these observations raise the possibility that some of these effects is through post-transcriptional mechanisms, such as mRNA stabilisation or miRNA sponging, rather than classical nuclear transcriptional activation.

Therefore, these findings may support a broader model in which cytoplasmic functions may contribute significantly to gene regulation and function. However, a comprehensive understanding of their subcellular distribution remains incomplete, and the functional significance of eRNAs beyond the nuclear compartment, particularly in the cytoplasm, has yet to be systematically explored. A centralised and accessible resource cataloguing eRNA localisation across subcellular compartments and mapping potential eRNA-eRNA interactions is currently lacking. This gap hampers efforts to distinguish nuclear-restricted enhancer activity from putative post-transcriptional functions and limits our ability to interrogate the full functional repertoire of eRNAs in gene expression control.

To address this gap, we developed **eRNAkit**, a comprehensive and user-friendly resource for exploring eRNA subcellular localisation, expression, and cytoplasmic functions. eRNAkit integrates publicly available fractionation-based transcriptomic datasets to systematically profile eRNA distribution across nuclear and cytoplasmic compartments. It also incorporates expression data across major human organs and primary cell types, allowing users to assess the tissue and cell-type specificity of individual eRNAs. Additionally, eRNAkit includes experimentally derived RNA-RNA interaction data from technologies such as RIC-Seq, PARIS, and KARR-Seq, enabling users to explore putative eRNA-eRNA interactions grounded in physical proximity and hybridisation evidence. By combining localisation, expression, and interaction data, eRNAkit offers a powerful framework to interrogate the full regulatory potential of eRNAs beyond the classical enhancer–promoter paradigm.

## Results

### eRNAkit Overview

eRNAkit represents the first comprehensive resource specifically designed to investigate the cytoplasmic functions of eRNAs (Figure 1). It integrates a suite of analytical workflows for processing and analysing eRNA-related data, facilitating tasks such as data manipulation, statistical analysis, and visualisation. To assist users, eRNAkit provides a transcriptome-wide landscape of eRNA subcellular localisation, eRNA-mRNA interactions and eRNA-ribosome associations, thereby elucidating their function beyond the nucleus. Compared to existing resources, eRNAkit offers the most extensive collection of cytoplasmic eRNA-specific functional information and analytical methods (Table 1), enabling a comprehensive interrogation of eRNA functions. To facilitate exploration of the database, it includes a user-friendly Rshiny interface.

**Figure 1:**
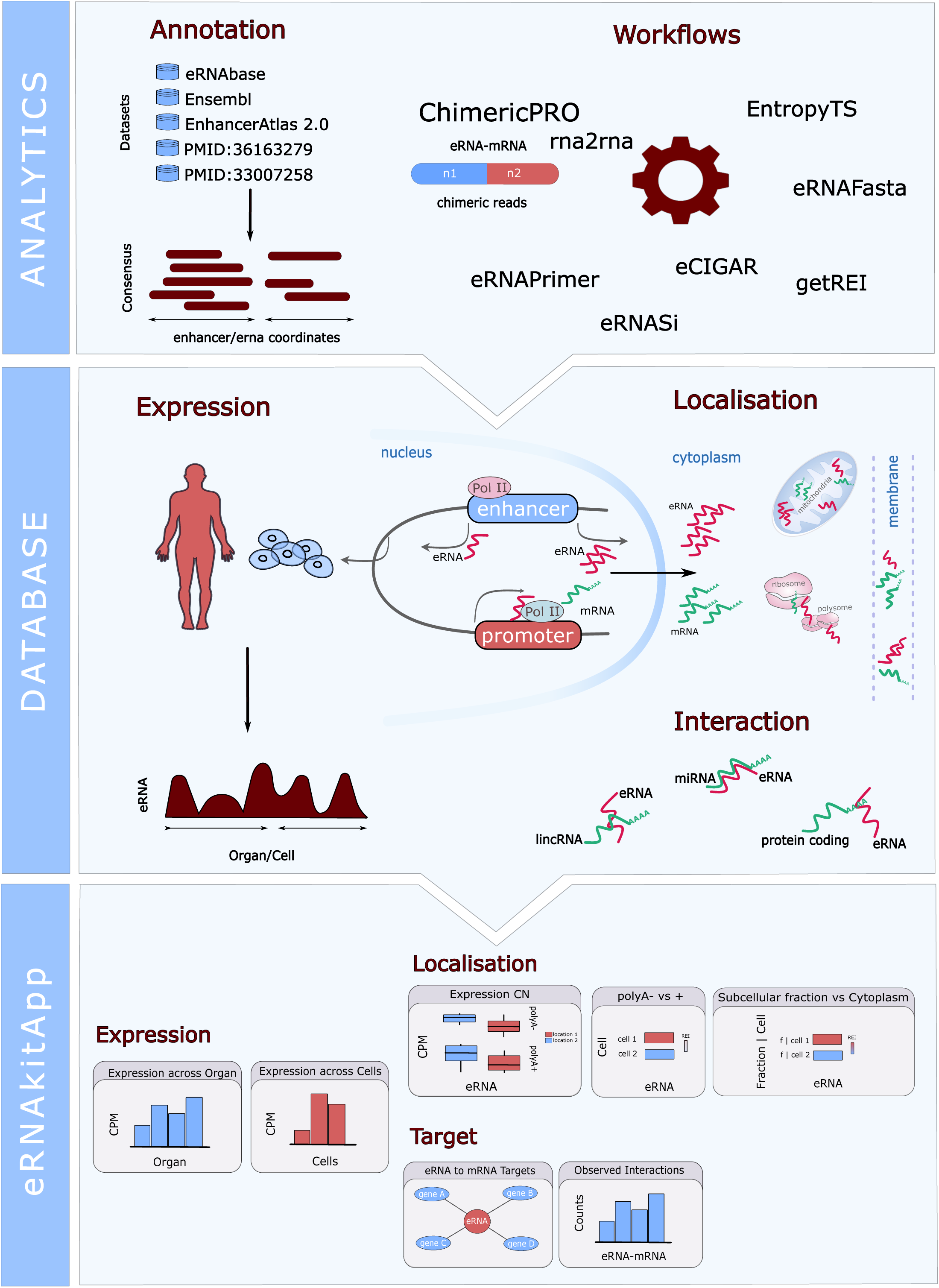
Overview of the main components of eRNAkit. eRNAkit consists of three layers: **Analytics**, **Database**, and **eRNAkitApp**. The **Analytics** layer provides workflows for processing eRNA-related data and supports the design of validation experiments. The **Database** contains data on eRNA expression across major tissues and primary cells, subcellular localisation, and physical eRNA–mRNA interactions. The **eRNAkitApp** offers a user-friendly interface for querying the database and includes modules for exploring eRNA function.

**Table 1:** Leading eRNA resources for comparison to eRNAkit.

|  | Subcellular localisation | mRNA interaction | Ribosome interaction | Target assignment |
| --- | --- | --- | --- | --- |
| **eRNAkit** | Nucleus, Cytosol, Nucleolus, Chromatin, Nucleoplasm, Membrane, Insoluble cytoplasmic fraction | KARR-Seq, RIC-Seq, PARIS | Ribo-Seq, Polysome-Seq | Physical eRNA-mRNA<br>(TAD used to differentiate nuclear from cytoplasmic) |
| eRNAbase | None | RIC-Seq | None | Genomic distance (multiple thresholds) |
| HeRA | None | None | None | Genomic distance (1MB) and Correlation ($\text{Rho} > 0.3$) |
| Anima — eRNAdb | None | None | None | Genomic distance (1MB) and Correlation ($\text{Rho} > 0.3$) |
| eRic | None | None | None | Genomic distance (1MB) |
| GPleR | None | None | None | Genomic distance (1MB) |

### eRNAkitApp

Extracting relevant information for a few eRNAs from the transcriptome-wide database can be tedious due to the complexity of navigating multiple interlinked tables. Thus, we build an interactive Rshiny UI called eRNAkitApp. The interface includes two core components: search panel and interactive visualisation modules. The search panel allows users to query the database by eRNA ID, genomic coordinates, or Ensemble gene ID, returning all relevant records. The visualisation modules include subcellular localisation (Supplementary Figure S1b), tissue/cell expression and specificity (Supplementary Figure S1c), eRNA-mRNA interactions (Supplementary Figure S1a) and eRNA-ribosome associations. Finally, the application offers straightforward results download option, allowing users to export search results tables. Additionally, the full database is also available for download for integration into other workflows. Collectively, this tool serves as a powerful platform for researchers to interactively explore and analyse eRNA-related data.

### Data Summary

We obtained 46,650 consensus eRNAs by integrating enhancer annotations from five large databases (Methods, Figure 1a). The median length of these eRNAs was 992 bp, with an interquartile range of 238–3,241 bp, and the longest extending up to 3,198,781 bp. We found 11 extremely large regions (> 300, 000 bp), which are derived from overlapping super-enhancers in the source databases. Across tissues and primary cells, 81.19% of the eRNAs were detected at a >1 CPM threshold (Supplemental Figure 1a-b). The number of detectable eRNAs ranged from 5,209 in the liver to 13,444 in the spinal cord for tissues, and from 1,957 in astrocytes to 20,688 in bladder smooth muscle cells (SMCs) for primary cells. Notably, SMCs and fibroblast populations accounted for the majority of the top 10 cell types. All tissues and cell types had at least one specific eRNA, with the vagina showing the highest number among organs (2,383) and mononuclear cells having the most among cell types (1,202).

Unlike protein-coding genes, the functional output of eRNAs is the RNA molecule itself. As such, their regulatory potential is closely linked to their subcellular localisation and physical interactions with other RNAs. Defining these spatial and molecular contexts is essential for understanding the mechanisms by which eRNAs exert their functions. As expected, more eRNAs are detectable in the nucleus, regardless of polyadenylation status (Figure 2b). However, polyadenylation influences the number of eRNAs detectable in the cytoplasm, potentially through controlling eRNA stability and transport. Analysis of the cytoplasm-nucleus REI distribution across cell types and RNA classes reveals a trimodal pattern, with subsets of eRNAs enriched in the nucleus, in the cytoplasm, or showing no strong bias (Figure 2c). While nuclear enrichment is more common, a substantial proportion of eRNAs are cytoplasm enriched. In the two cell lines with subcytoplasmic fractionation data, we observe widespread localisation of eRNAs in both the insoluble (organelle-associated) and membrane fractions (Figure 2d).

**Figure 2:**
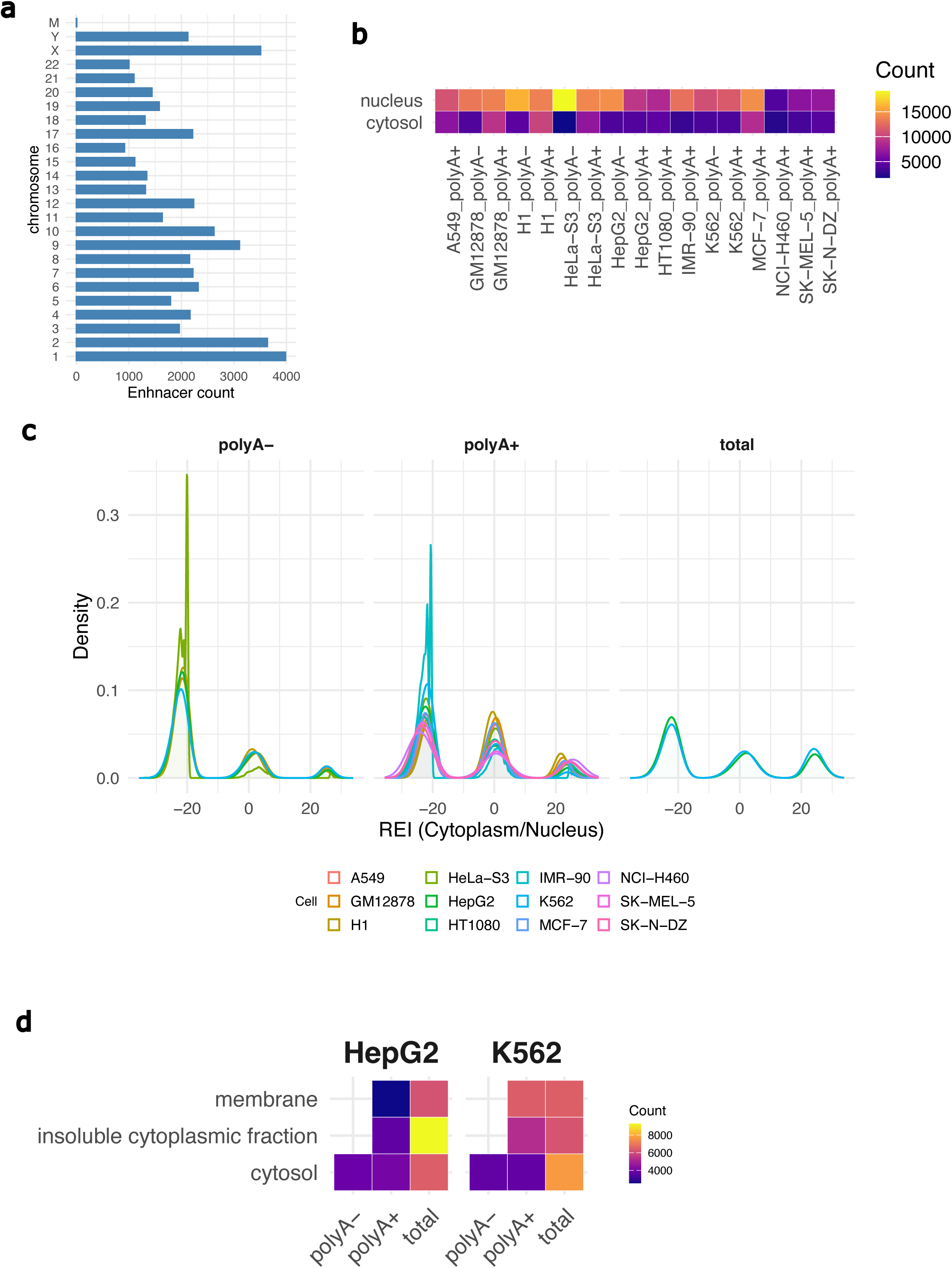
Summary statistics from the database layer. **(a)** Distribution of consensus eRNAs across chromosomes. **(b)** Number of detected eRNAs (CPM > 1) in both the nucleus and cytoplasm across multiple cell types. polyA+ indicates selection of polyadenylated RNAs prior to sequencing, while polyA− indicates depletion of polyadenylated RNAs. **(c)** Distribution of the relative expression index (REI) between cytoplasm (C1) and nucleus (C2) across RNA classes (see Methods). Positive values indicate enrichment in the cytoplasm, and negative values indicate enrichment in the nucleus. **(d)** Number of detected eRNAs (CPM > 1) in cytoplasmic sub-fractions across RNA classes in HepG2 and K562 cells.

Our curated dataset of eRNA-mRNA interactions identified 82,587 pairs present in at least two samples, encompassing 25.28% (11,793 out of 46,650) of the annotated consensus eRNAs (Figure 3a). Given the conservative nature of our filtering, this number is expected to increase with broader dataset inclusion. Surprisingly, correlation analysis between eRNA length and the number of interactions revealed a significant weak negative correlation (*rho* = −0.207, *p* < 2.2 × 10⁻¹⁶; *S* = 3.30 × 10, Figure 2a). This suggests that longer eRNAs may be slightly less likely to engage in RNA-RNA interactions. However, the low correlation coefficient indicates that factors beyond transcript size-such as sequence features, localisation, or binding affinity-likely contribute to their interaction potential. Indeed, stratified analysis of these interactions across diverse cellular perturbations reveals that some are dependent on active translation or RNA-binding proteins (Figure 3b). Notably, similar perturbations-such as translation inhibition by cycloheximide or harringtonine, and viral infection by RSV or VSV-lead to consistent alterations in the RNA-RNA interaction landscape (Figure 3b). Additionally, only 28.7% (23, 703 out of 82, 587) of the mRNA were located within the same TAD or within 5kb of their corresponding eRNA pair (Methods). Interestingly, 30% of all the interactions pairs occur between eRNAs and mRNAs located on different chromosomes. These observations suggest that most eRNA-mRNA interactions do not adhere to the canonical *cis*-regulatory function of eRNAs.

**Figure 3:**
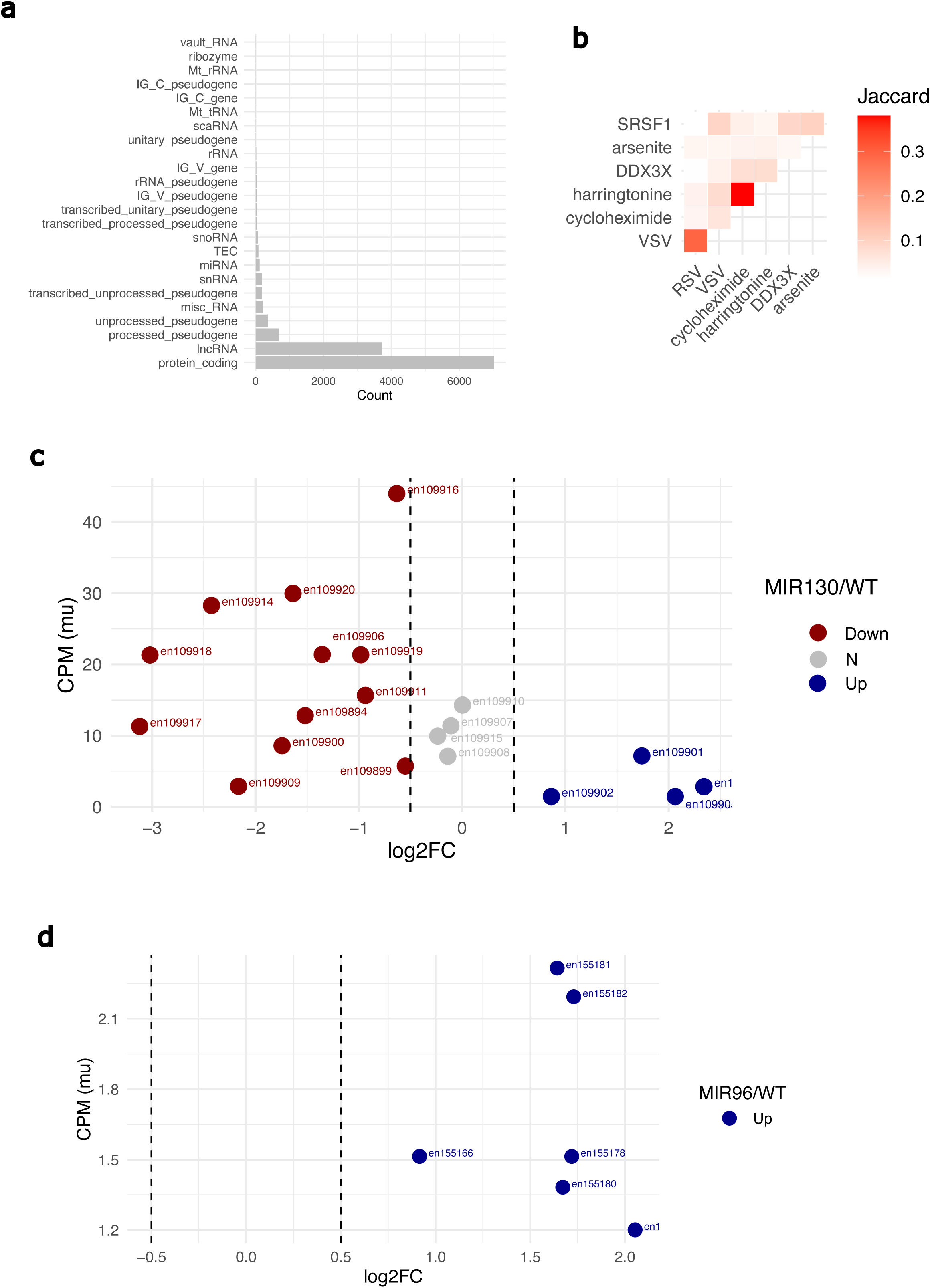
eRNA–mRNA interactions reveal potential cytoplasmic functions. **(a)** Distribution of eRNA–mRNA interactions across mRNA biotypes. **(b)** Similarity of eRNA–mRNA interactions (gain or loss) under different cellular perturbations. SRSF1 and DDX3X represent knockdowns of RNA-binding proteins. Harringtonine and cycloheximide indicate translational inhibition via drug treatment. RSV and VSV represent infection with respiratory syncytial virus or vesicular stomatitis virus, respectively. Arsenite denotes induction of cellular stress. **(c–d)** Scatter plots showing the median expression of eRNAs in control cells versus the log₂ fold change in expression upon perturbation with miR-130 **(c)** or miR-96 **(d)**.

### Case study

#### Potential miRNA sponging by eRNAs

An established cytoplasmic function of several classes of non-coding RNAs is the ability to act as miRNA sponges — sequestering miRNAs and preventing them from repressing their target mRNAs. This mechanism has been widely described for circular RNAs and long non-coding RNAs ^14,15^. Given this context, it is plausible that eRNAs, especially those with stable cytoplasmic presence — may also function as miRNA sponges. This possibility has not been systematically explored, but the pervasive localisation of eRNAs hints at such roles. To illustrate this, we examined the eRNA targets of miR130b and miR96, both of which have been implicated in diverse cellular processes, including metabolism and tissue development. miR-130B has been shown to regulate adipogenesis ^16^, while miR-96 is involved in neuronal cell differentiation ^17^. Using eRNAkitApp, we first input their Ensemble IDs “ENSG00000283871, ENSG00000199158” and selected **TARGET** (Figure S1a). The results showed two distinct clusters of target eRNAs, indicating potential differences in function. Then we selected the **LOCALISATION** tab, which revealed that most of the associated eRNAs are detectable in the cytoplasm and sub-cytoplasmic fractions (Figure S1b). Given the strong cytoplasmic localisation of these eRNAs in this context, and the precedence of miRNA sponging among other cytoplasmic non-coding RNAs, we proposed that miRNA sequestration may be a functional mechanism for these eRNAs. To investigate this possibility, we examined changes in eRNA expression in response to miR-130 overexpression in mesenchymal stem cells (GSE202135) and miR-96 knockdown in prostate epithelial cells (GSE255227). Following miR-130 overexpression (by mimics), 60% (12 out of 20) of the detectable (CPM > 1 in > 2 samples) eRNA targets were downregulated (log2FC < - 0.5), a proportion markedly higher than expected by chance (empirical p = 0.0, Figure 3c). By contrast, all detectable eRNA targets of miR-96 (6 out of 6) were upregulated (log2FC > 0.5) upon miR-96 knockdown (by siRNA), also surpassing random expectation (empirical p = 0.0019, Figure 3d). These findings suggest that, like other cytoplasmic ncRNAs, eRNAs may contribute to post-transcriptional regulation by modulating miRNA availability and illustrate the biological value of the eRNAkit.

#### Inter-organelle communication through eRNAs

Bi-directional communication between the nucleus and mitochondria is critical for maintaining cellular function ^18^. Emerging research suggests that RNA-mediated nuclear signal can influence mitochondrial biology ^19^. We identified an eRNA, “en135763” located on chromosome 5, which is altered under various cellular stress conditions (data not shown), and searched eRNAkitApp using the eRNA ID option (Figure S3a). In the **LOCALISATION** tab, results showed that this eRNA exhibits comparable expression levels in the cytoplasm and nucleus. It is also enriched in the organelle-associated fraction in K562 cells and has eRNA-RNA interactions with mitochondrial encoded genes (Figure S3b), indicating that eRNAkit could support research into RNA mediated inter-organelle communication. The **EXPRESSION** tab revealed that this eRNA is widely expressed across organs and primary cell types, with notably higher levels in T-cell populations, consistent with the involvement of immune cells in the stress response. These findings highlight the underappreciated value of eRNA subcellular localisation and physical interactions.

## Discussion

eRNAs have long been considered nuclear-localised and unstable byproducts of enhancer activation, with functions largely confined to cis-regulatory roles at nearby promoters. However, emerging data challenge this restricted view, revealing that eRNAs are more biochemically and functionally diverse than previously appreciated. Unlike protein-coding genes, the final product of eRNAs is RNA itself. Consequently, eRNA function relies on the physical properties and interactions of the RNA molecule, which are inherently dependent on spatial proximity. Here, we present eRNAkit, a comprehensive resource that consolidates subcellular localisation, expression specificity, and RNA-RNA interaction data to systematically map the full regulatory repertoire of eRNAs, including potential trans-acting and post-transcriptional functions. Compared to existing databases, eRNAkit offers several advances. Most notably, it is the first to integrate larger scale subcellular localisation data alongside physical RNA-RNA interactions, thereby facilitating functional hypotheses that extend beyond proximity-based enhancer-promoter models.

A key component of eRNAkit is the transcriptome-wide landscape of eRNA localisation across subcellular compartments. Our REI-based analyses demonstrate that while nuclear enrichment is prevalent, a considerable proportion of eRNAs are detectable — and in some cases, enriched in cytoplasmic and subcytoplasmic fractions. This observation supports prior reports suggesting the presence of cytoplasmic eRNAs ^3^ and offers a transcriptome-wide perspective that challenges the long-standing assumption of nuclear restriction. Notably, the influence of polyadenylation on cytoplasmic localisation suggests that RNA processing features may be critical determinants of eRNA stability and trafficking, potentially mirroring the behaviour of other non-coding RNAs with post-transcriptional roles, such as lncRNAs and circRNAs. eRNAkit also highlights strong cell- and tissue-specific expression patterns of eRNAs, consistent with the known context-dependence of enhancer activity ^1,20^. The presence of highly specific eRNAs in reproductive organs and in immune cells highlight their potential roles in tissue-specialised gene regulatory programs. Such specificity may be leveraged in future studies to prioritise candidate eRNAs for functional validation or biomarker development.

The inclusion of tens of thousands of robust eRNA-mRNA interactions further expands the functional landscape of eRNAs. These interactions, derived from multiple RNA-RNA interactome technologies (RIC-Seq, PARIS, KARR-Seq), provide direct evidence of physical proximity between eRNAs and potential target transcripts. Importantly, a substantial fraction of these interactions is translation or RBP-dependent, suggesting that eRNAs may engage in post-transcriptional regulation through mechanisms such as mRNA stabilisation, translational control, or miRNA competition. The weak negative correlation between eRNA length and interaction count also points to a nuanced interaction landscape that may be shaped more by structure, binding motifs, or localisation than by transcript size alone.

Despite its utility, several limitations should be noted. First, while the use of multiple datasets enhances confidence in reported interactions and expression patterns, coverage across all cell types and compartments remains incomplete. Second, although physical interactions provide strong evidence for potential function, mechanistic validation is essential to determine whether these interactions are regulatory, structural, or stochastic. Lastly, the dynamic regulation of eRNA expression and localisation — particularly under stress, development, or disease contexts — remains to be fully characterised and could provide further insight into the functional adaptability of eRNAs. To address these limitations, we plan to update eRNAkit regularly by incorporating additional subcellular localisation data and functional eRNA interaction profiles as more high-throughput datasets become available. Due to our work in cellular senescence, we are also building an extension to characterise the expression profiles of these eRNAs in cellular stress response. We are confident that eRNAkit provides a critical first step toward uncovering the full regulatory potential of eRNAs and will serve as a foundational tool for advancing our understanding of their diverse roles in human health and disease.

## Methods

### eRNAkit Design

eRNAkit comprises three main layers: analytics, database and Rshiny application (Figure 1). The analytics layer includes a suite of functions designed to characterise the cytoplasmic functions of eRNAs. Key routines encompass rna2rna for identifying eRNA-mRNA interactions from STAR aligner chimeric junctions, entropyTS for assessing tissue or cell specificity and getREI for calculating the relative expression index. The database layer, referred to as emi, contains four structured tables that store consensus eRNA annotations, transcriptome-wide eRNA-mRNA interactions, subcellular localisation data of eRNA classes and their association with ribosomes, as well as expression levels of eRNAs across major human organs and primary cell types, including tissue and cell-type specificity metrics. Additionally, the database integrates genome-wide enhancer-target gene information, facilitating the differentiation between nuclear and cytoplasmic functions of eRNAs. The R Shiny application layer provides an interactive interface for users to explore and visualise the database, enabling dynamic querying and presentation of results.

### Consensus human eRNA annotation

To build consensus eRNAs annotation, we collected previously published human enhancer elements across multiple tissues and cell types from EnhancerAtlas 2.0 ^21^, Ensemble regulatory build ^22^, eRNAbase ^5^, neuronal transcribed enhancers ^23^, and cancer associated eRNAs ^24^. We manually downloaded 2, 492, 442 coordinates in bed format and converted those in hg19 genome build to hg38 coordinates using liftOver tool ^25^. Then, overlapping elements across the five datasets were merged to generate a consensus file using BEDTools^26^. To ensure robust genomic coordinates, we validated all annotations against the hg38 chromosome sizes, excluding any entries with start or end positions outside of chromosome bounds. To ensure that eRNA quantification is not confounded by normal gene expression, we removed elements overlapping with the Ensemble gene annotation v112. We extended the gene regions by 500 bp at both sides for this analysis to remove possible unknown variants. Finally, we removed elements with a size < 10 bp, which are within the range of largely unstable nuclear eRNAs.

### eRNA subcellular localisation

Subcellular RNA-Seq data representing two RNA classes (polyA− and polyA+) across seven cellular fractions (nucleus, cytosol, nucleolus, chromatin, nucleoplasm, membrane, insoluble cytoplasmic fraction) were obtained from ENCODE ^3^. BAM files aligned to the hg38 human genome assembly for 13 cell lines (SK-N-SH, K562, A549, H1, HeLa-S3, MCF-7, HepG2, IMR-90, GM12878, SK-N-DZ, SK-MEL-5, NCI-H460, and HT1080) were downloaded from the ENCODE Data portal (Supplemental Table S2). Read counts were quantified using HTSeq (v0.11.1)^27^ on the consensus eRNA GTF file. Each RNA class and subcellular fraction had biological replicates, and read counts were summarised by the median count per million (CPM) across replicates.

To assess relative compartment expression of eRNAs across RNA classes and cell types, we calculated the relative expression index (REI) across predefined pairs of subcellular compartments for each RNA class (polyA−, polyA+ or total RNA) and cell line. REI was defined as:

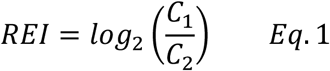

where *C_1_* and *C_2_* represent the median of a given eRNA in two compartments. A higher REI value indicates greater expression in *C_1_* relative to *C_2_*.

Compartment pairs were defined such that finer cytoplasmic subfractions (e.g., insoluble cytoplasmic fraction, membrane) were compared to the cytosol, and nuclear subfractions (e.g., nucleoplasm, nucleolus, chromatin) were compared to the nucleus. The complete set of compartment comparisons were cytosol-nucleus, insoluble cytoplasmic fraction-cytosol, membrane-cytosol, nucleolus-nucleus, nucleoplasm-nucleus, chromatin-nucleus. For each RNA class and cell line, REIs were computed only when expression values were available for at least one compartments being compared. To investigate eRNA processing, a second REI was calculated between polyA− and polyA+ samples within each compartment and cell using the same filtering strategy.

### Calculation of tissue and cell specificity for eRNAs

To assess tissue and cell specificity, we downloaded and processed total RNA-Seq BAM files (hg38) for 32 major tissues and 77 primary cell types from the ENCODE data portal. Biological replicates were summarised by the median CPM across replicates. eRNAs that were lowly expressed (<1 CPM) in all tissues were removed. We call an eRNA specific (tissue or cell) when its maximum median CPM was more than double the second largest median CPM and the difference between the logarithm of the total number of tissues/cells and the Shannon entropy of the median CPMs is greater that 1 ^28^.

### eRNA-mRNA interaction

The eRNA-mRNA interactions in eRNAkit were identified using three high-throughput approaches, including KARR-Seq ^29^, RIC-Seq ^30^ and PARIS ^31^ (Table 2). These methods differ in their underlying chemistries, interaction specificity, and compartmental bias, together providing a more comprehensive and nuanced map of eRNA-mRNA contacts. By integrating datasets with differing sensitivities to nuclear versus cytoplasmic interactions, eRNAkit enables exploration of potential post-transcriptional regulatory roles for eRNAs that might be overlooked by any single approach.

**Table 2:** Comparison of RNA–RNA Interaction Mapping Methods.

| Feature | KARR-Seq | RIC-Seq | PARIS |
| --- | --- | --- | --- |
| **Chemistry** | Kethoxal-modified guanine bases (selective to ssRNA) | Formaldehyde (reversible; targets protein and RNA interactions) | Psoralen (intercalates in double-stranded RNA) |
| **Interaction** | Base-paired duplexes, especially transient or regulatory pairs | Both direct and indirect (via RNA-binding proteins) interactions | Base-paired duplexes with high confidence |
| **Specificity** | Prefers G-rich and dynamic regions | Protein-mediated proximity bias | Enriches for stable RNA duplexes |
| **Compartment Bias (Whole Cell Lysate)** | **Cytoplasmic-enriched** — favours mature, exported RNAs | **Balanced** — captures both nuclear and cytoplasmic interactions | **Nuclear-biased** — stronger detection of pre-mRNA and nuclear lncRNA |
| **Advantages** | Sensitive to dynamic, potentially regulatory interactions | Captures a broad spectrum of RNA interactions | Provides structural information; precise mapping of duplexes |
| **Limitations** | May miss highly structured or non-G-rich regions | Less resolution for direct base-pairing | Bias toward stable, long helices |

We manually downloaded 78 raw sequencing datasets from 9 cells lines via the NCBI GEO database (Supplemental Table S1). To identify eRNA-mRNA interactions, reads were quality filtered (Q < 20) and adapter trimmed using Trimmomatic (v0.39) ^32^. Cleaned reads were aligned to the human reference genome (hg38) using STAR ^33^ with chimeric alignment options enabled (--runMode alignReads --chimOutType Junctions --chimJunctionOverhangMin 8 --chimSegmentMin 15 --outFilterMultimapNmax 100 --alignIntronMin 1 --scoreGapNoncan −4 --scoreGapATAC −4 --alignSJoverhangMin 15 --alignSJDBoverhangMin 10 --alignSJstitchMismatchNmax 5 −1 5 5).

Chimeric junctions were then processed using a custom R-based workflow implemented in eRNAkit (rna2rna). Briefly, chimeric alignments were parsed into a pseudo-BEDPE format by splitting each read into two genomic intervals (n1 and n2) and extracting positions and strand information using the CIGAR string. These were intersected with Ensemble v111 gene annotations and the consensus eRNA annotations. Only junctions connecting an eRNA to a known gene were retained as putative eRNA-mRNA interactions. To enhance robustness, only eRNA-mRNA pairs detected in at least two independent samples were included in the final interaction set. This conservative filtering strategy prioritises reproducibility across biological contexts and offers a high-confidence, global view of eRNA interactions.

Some of the samples analysed by KARR-Seq included cellular perturbations such as translation inhibition, RNA-binding protein knockdowns and viral infections (Supplemental Table S1). These experimental contexts provided an opportunity to explore potential mechanisms driving eRNA-mRNA interactions. To assess perturbation responsiveness, we focused on binary changes-specifically, the complete gain or loss of individual interactions in treated versus untreated conditions. Interactions were considered perturbation-sensitive if they were entirely absent in control samples but appeared in treated samples, or vice versa. This stringent approach highlights interactions potentially under regulatory control in response to environmental or cellular cues.

### eRNA genomic targets

To identify potential genomic targets to facilitate differentiation of cytoplasm exclusive interactions, we used two strategies (1) distal targets, based on co-localisation within the same topologically associating domains (TADs), and (2) proximal targets, based on linear distance to the eRNA loci. For the distal approach, we obtained human TAD annotations across multiple cell types and tissues from the ENCODE database ^34^. Using BEDTools (v2.31.1) ^26^ and human gene annotation (hg38, Ensemble v111), we identified genes residing in the same TAD as each eRNA locus. For the proximal approach, genes located within ±5 kb of an eRNA locus were annotated as their proximal targets. Note that both approaches mirror what is used by existing tools to call eRNA targets (Table 1).

### RNA-Seq and Ribo-Seq analysis

Standard RNA-Seq and Ribo-Seq (Supplemental Table S3) analysis was performed using our existing pipeline ^35^. Briefly, the raw reads were quality filtered (Q < 20) and adapter trimmed using Trimmomatic (v0.39) ^32^. Cleaned reads were aligned to the human reference genome (hg38) using HISAT2 (v2.1.0) in default settings ^36^. Read counts were quantified using HTSeq (v0.11.1)^27^ on the consensus eRNA annotation. The expression levels were normalised by CPM.

### Permutation test

To determine whether the number of differentially expressed targets associated with a given miRNA was significantly greater than expected by chance, we performed a non-parametric permutation test. The observed count was defined as the number of eRNA targets in a specific threshold (Log_2_ fold change (Log2FC) > 1 or < −1) among the set of detected eRNA targets of miR-130b or miR-96. To generate the null distribution, we randomly sampled the same number of eRNA (x10,000) from all the detected eRNA and counted the number matching the threshold. The empirical p-value was computed as the proportion of permutations in which the number of random eRNAs in the specified threshold was greater than or equal to the observed count. This one-tailed test estimates the probability of obtaining the observed level of enrichment under the null hypothesis of random association.

## Data availability

The research community can access eRNAkit freely from the GitHub repository https://github.com/AneneLab/eRNAkit.

## Author contributions

Project was conceived of by C.A.A. Data curation and processing were done by N.B, R.K, M.E, and C.A.A. Software and database creation was done by C.A.A, N.B and R.K. C.A.A supervised the project and wrote the manuscript. All authors edited the final manuscript.

---

**Figure S1:**
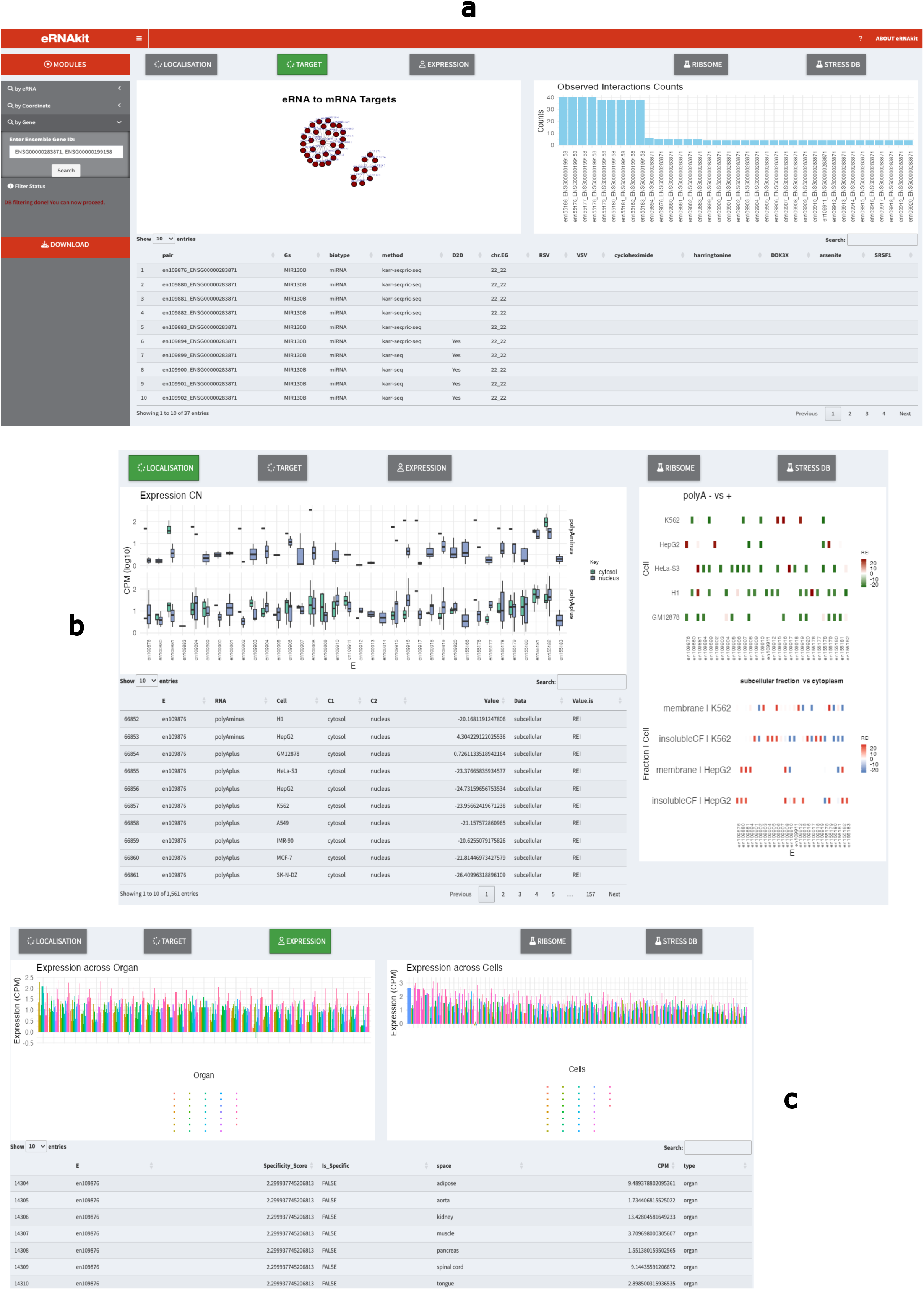
Main interface and usage of eRNAkitApp. **(a)** Navigation bar with the TARGET module activated. **(b)** View of the LOCALISATION module. **(c)** View of the EXPRESSION module. **Note:** Graphs shown in S1 relate to searching for “ENSG00000283871, ENSG00000199158” (MIR130 and MIR96).

**Figure S2:**
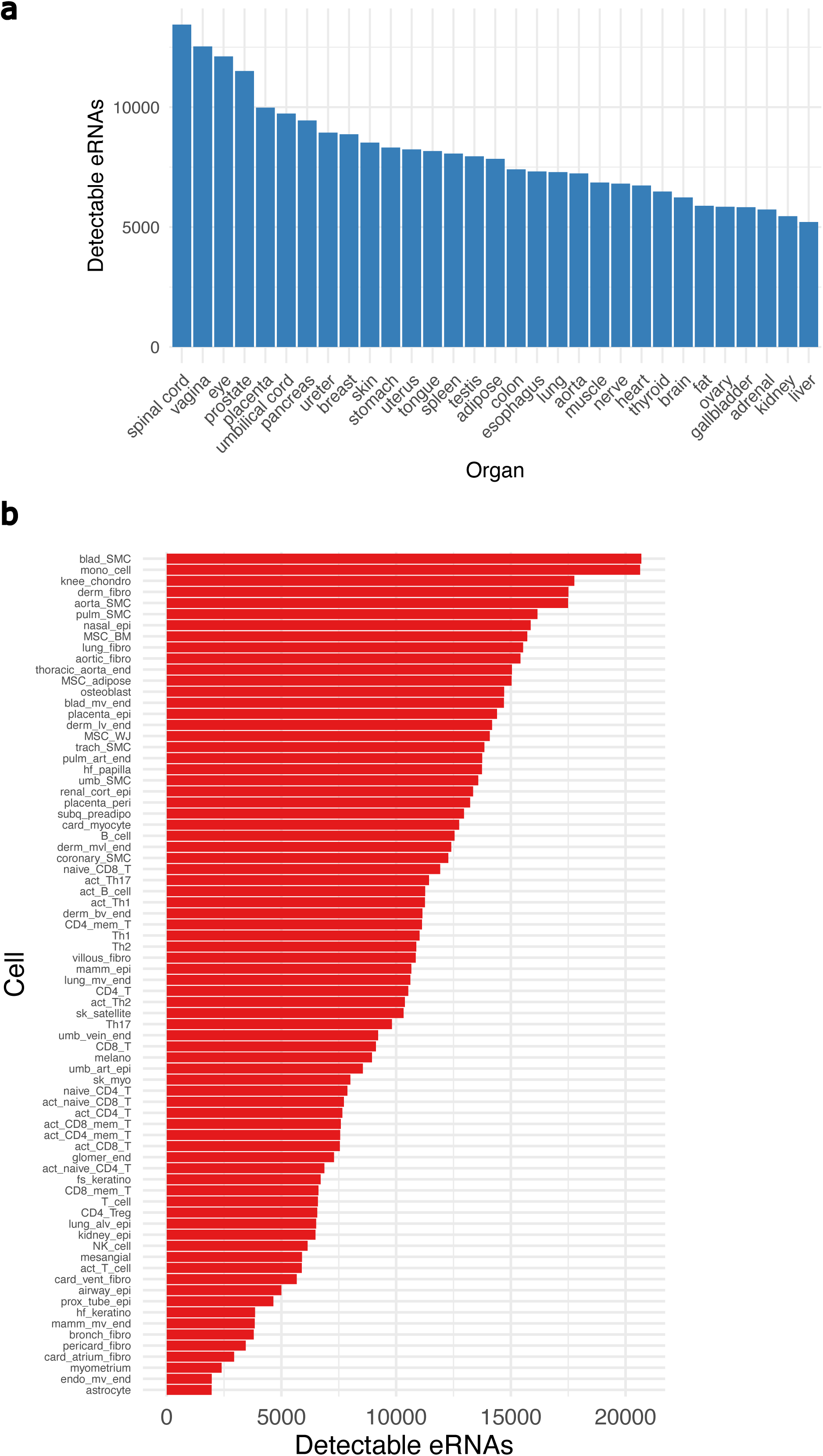
eRNAs are widely expressed across major tissues and primary cells. **(a–b)** Number of detected eRNAs (CPM > 1) across tissues **(a)** and primary cells **(b)**.

**Figure S3:**
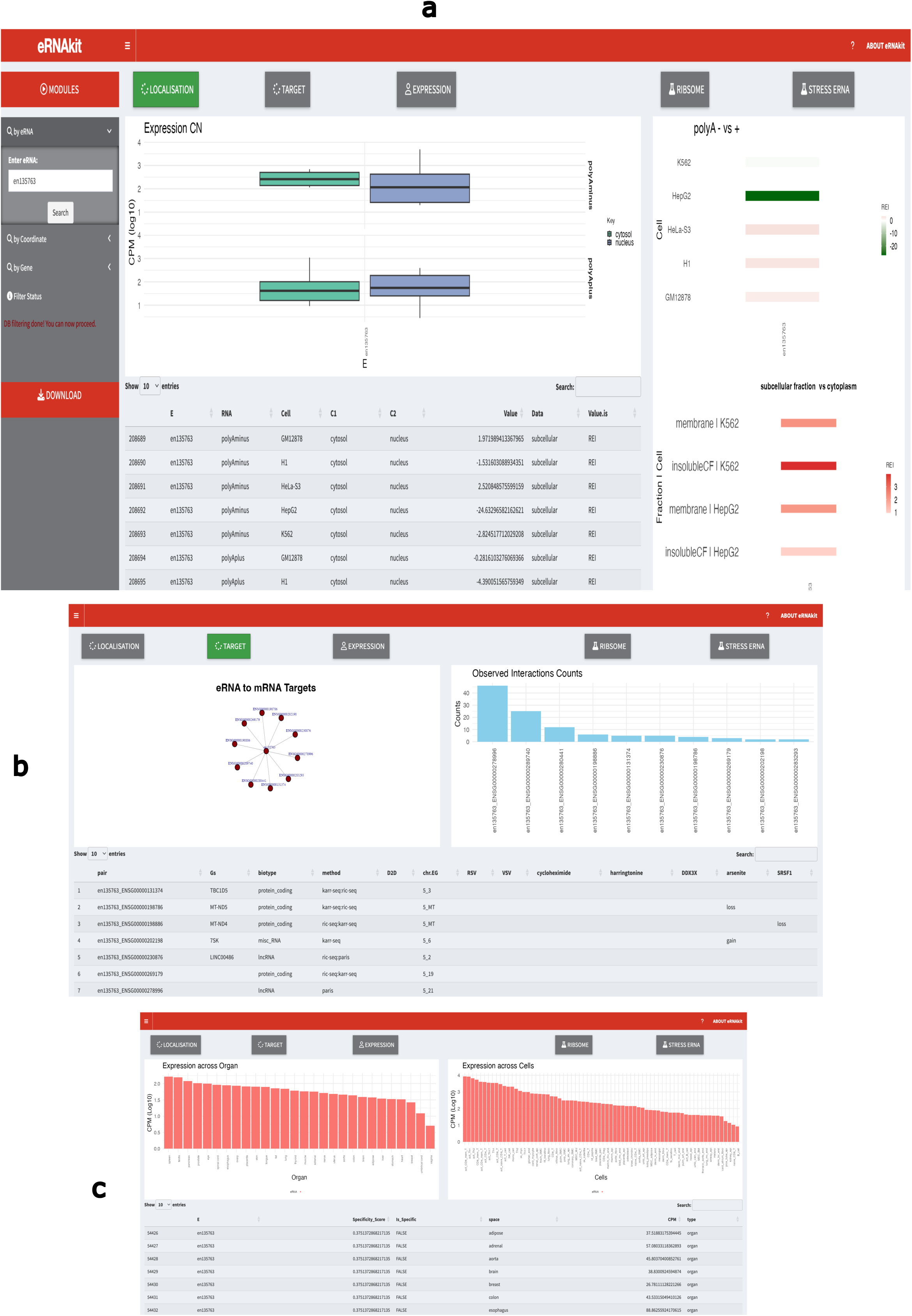
Searching eRNAkitApp using eRNA ID. **(a)** Navigation bar with the LOCALISATION module activated. **(b)** View of the TARGET module. **(c)** View of the EXPRESSION module. **Note:** Graphs shown in S3 relate to searching for “en135763”.

